# CASPAM: a triple modality biosensor for multiplexed imaging of caspase network activity

**DOI:** 10.1101/2021.03.10.434623

**Authors:** Martín Habif, Agustín A. Corbat, Mauro Silberberg, Hernán E. Grecco

## Abstract

Understanding signal propagation across biological networks requires to simultaneously monitor the dynamics of several nodes to uncover correlations masked by inherent intercellular variability. To monitor the enzymatic activity of more than two components over short time scales has proven challenging. Exploiting the narrow spectral width of homoFRET-based biosensors, up to three activities can be imaged through fluorescence polarization anisotropy microscopy. We introduce CASPAM (Caspase Activity Sensor by Polarization Anisotropy Multiplexing) a single-plasmid triple-modality-reporter of key nodes of the apoptotic network. Apoptosis provides an ideal molecular framework to study interactions between its three composing pathways (intrinsic, extrinsic and effector). We characterized the biosensor performance and demonstrated the advantages that equimolar expression has both in simplifying experimental procedure and reducing observable variation, thus enabling robust data-driven modelling. Tools like CASPAM become essential to analyze molecular pathways where multiple nodes need to be simultaneously monitored.

## Introduction

Molecular signaling networks act together to control cellular dynamics. The complex interactions between different intracellular pathways determine their spatiotemporal coordination and ultimately define the biological functionality [1, 2]. Hence, comprehensive understanding of the hierarchy and cross-regulatory linkages between the distinct molecular pathways becomes essential.

Fluorescence microscopy methods provide currently the only tools to examine the dynamic properties of living systems with high temporal and spatial resolution [3]. As multiplexed imaging can reveal the coordination of both manifold subcellular structures and protein activities [4, 5, 6] there is a continuous motivation to develop innovative fluorescence-based biosensors to fulfill this goal [7]. However, connections between different nodes in a network are often inferred indirectly. In this sense, one of the biosensors may itself be used as the common reference in a series of separated experiments, where it is visualized simultaneously in combination with other reporters [6, 8]. Pathway reconstruction is then achieved by using the so-called *computational multiplexing* procedure, which enables for the correlation of many cellular signaling activities based on statistical approaches [6, 9].

Genetically encoded hetero-FRET-based biosensors (consisting of two spectrally separated fluorescent proteins (FP): a donor and a red-shifted acceptor) have uncovered relevant spatiotemporal aspects of protein interaction networks [10, 11]. Despite this, due to the broad spectra of FP, co-imaging in the visible range is limited to two sensors (4 FP) [1, 12]. In this sense, single fluorophore (or spectrally similar fluorophore) homo-FRET based biosensors considerably shrink the spectral window per sensor by means of fluorescence polarization anisotropy (FPA) [13, 14, 15], thus enabling the insertion of up to three reporters in a single cell [12].

Apoptosis provides the ideal molecular framework for studying the interactions between the multiple components of an intracellular signaling cascade. The machinery responsible for the apoptotic process relies on the sequential action of an evolutionarily conserved family of cysteine-proteases called caspases [16, 17]. Intrinsic apoptotic pathway is triggered by sources of stress originating within the cell, such as oncogenes, direct DNA damage and hypoxia [18]. Alternatively, extrinsic apoptosis involves extracellular ligand mediated activation by the so-called family of death-receptors [19]. Intrinsic and extrinsic apoptosis pathways imply the initial action of caspase 9 and 8, respectively, which finally conclude in the activation of effector caspase 3 [20]. Therefore, understanding signal propagation throughout the apoptotic network requires monitoring at least the activity of three different enzymes and, hence, the controlled insertion of three different biosensors in a single cell.

Co-expression of multiple biosensors has always been presented as a major experimental challenge. On one side, regardless of the carrier by which genetically encoded biosensors are delivered into cells, the success rate for their simultaneous incorporation is often sparse. Moreover, additional delays due to enzyme sequestration are introduced in signal propagation once the sensors are expressed. Different stoichiometric relationships between sensors probing distinct pathways within the network result in variability in introduced delays.

In this work, we report the development of CASPAM (**C**aspase **A**ctivity **S**ensor by **P**olarization **A**nisotropy **M**ultiplexing), a novel single-plasmid-based triple-modality reporter-system for simultaneously assessing the spatiotemporal activity dynamics of three different caspases, at the single cell level. CASPAM was validated in vitro in HeLa cells, and showed significantly improved performance over other available experimental alternatives in terms of co-expression ratio and stoichiometry. We were able to weigh up the advantages of CASPAM through its implementation in determining the activity of different signaling nodes in the apoptotic network. The use of CASPAM resulted both in the enablement of multiparametric single-cell analysis and the overcoming of unwanted intracellular perturbations due to heterogeneities in biosensor stoichiometry, which otherwise may affect the transmission and flow of information.

## Results

### Construction of CASPAM

Homo-FRET measured by fluorescence polarization anisotropy (FPA) microscopy has been used as a tool to detect molecular interactions, conformational changes and assembly state [12, 21, 14]. Briefly, the emission of FP is highly anisotropic upon excitation with linearly polarized light, due to the rather larger size and therefore their large rotational diffusion time as compared to the fluorescence lifetime. However, when another FP is in proximity, the emission becomes more isotropic due to homo-FRET. Therefore, two FP linked by a cleavage site serve as a reporter of protease activity, where its state can be inferred from the anisotropy of the emission [22, 23] (**Figure 1A**).

**Figure 1:**
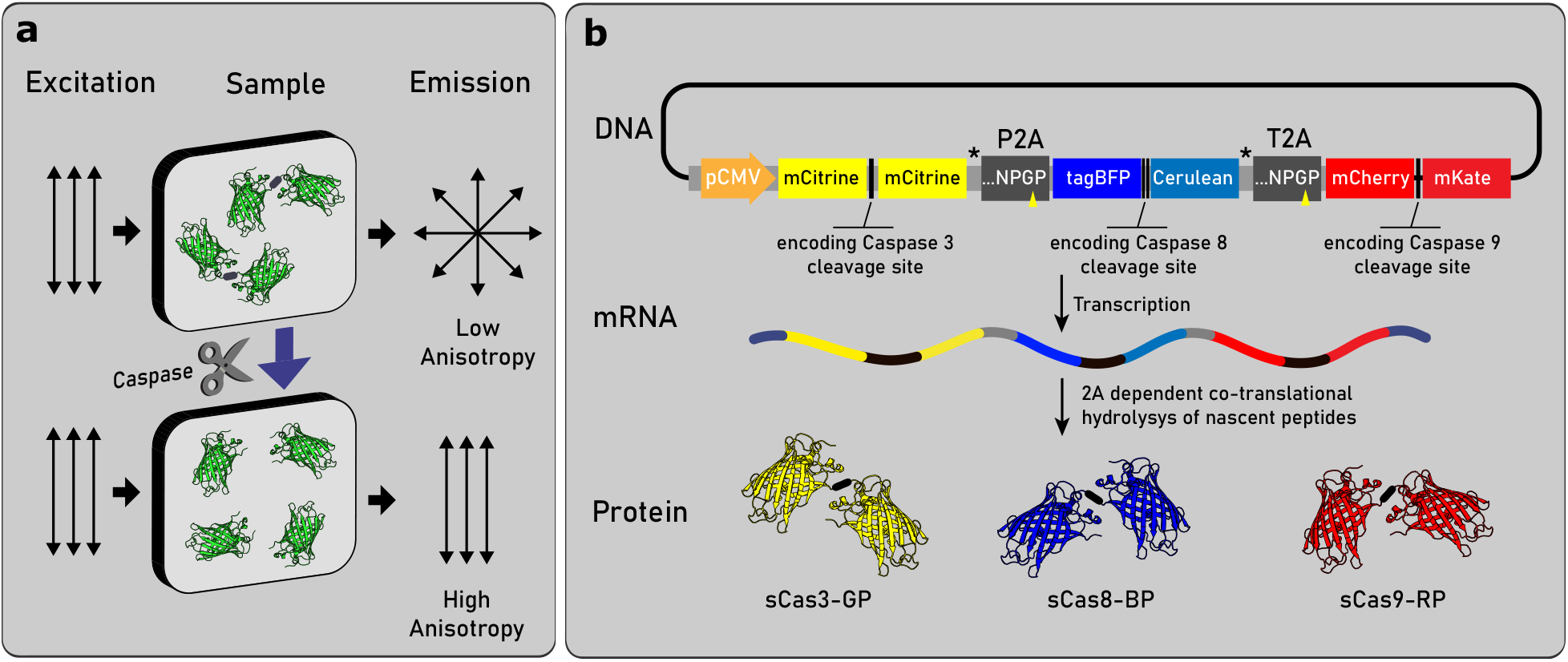
Spectrally separable anisotropy FRET-based sensors state can be determined by fluorescence polarization microscopy. **(a) Fluorescence Polarization Anisotropy (FPA) rationale**. Identical or spectrally similar FP within a sensor are linked by a sequence specifically recognized by the protease of interest. Following excitation with linearly polarized light, the emission of the ensemble of FP exhibits low polarization anisotropy due to the occurrence of FRET in dimers (top scheme). Conversely, monomeric state resulting from enzyme-mediated dimer cleavage render a more polarized emission (bottom scheme). As the reaction is unidirectional, the fraction of cleaved sensor is therefore a measure of the integral activity of the protease. **(b) CASPAM design rationale**. Diagram of the mammalian expression plasmid harboring a tandem array of the coding sequences for the three homo-FRET caspase activity sensors with viral 2A cassettes in between. DNA sequences encoding the cleavage sites for caspase 3, 8 and 9 (DEVD, IETD-IETD, LEHD, respectively) were located between the contiguous fluorescent ORFs of each biosensor. After CASPAM is transcribed as a single mRNA, 2A dependent hydrolysis occurs co-translationally, resulting in three independent biosensors. Flexible GGGSGGG linkers [24] were added to improve hydrolysis efficiency [39, 40, 25]

As previously described for multiprotein expression arrangements, where co-expression from a single ORF is successfully achieved [24, 25, 26, 27], we made use of 2A-peptide cis-acting hydrolase elements to co-express FPA caspase activity sensors at close to equimolar amounts. Therefore, we engineered CASPAM as a single mammalian expression plasmid harboring a tandem array of the sequences encoding three homo-FRET caspase activity sensors, with the viral 2A sequences in between.

For the sake of simplicity, we are going to name the biosensors according to the caspase sensed and the color of the fluorophore-pair used. In this way, tagBFP-x-Cerulean, mCitrine-x-mCitrine and mCherry-x-mKate will be named sCasx-BP, sCasx-GP and sCasx-RP. Three different target sequences for caspase 3, 8 or 9 specific cleavage were implemented: DEVD, IETD-IETD and LEHD, respectively. The coding sequences included in the construct (**Supplementary Figure S1**), that altogether lead to the synthesis of an expected *∼*5Kb mRNA, were under the control of the single CMV promoter contained in a pcDNA3.1(+) vector. As comparative studies show that self-cleavage efficiency decreases among existing 2A sequences in the order P2A*<*T2A*<*E2A*<*F2A [28, 29], we used for our multicistronic expression system, 2A peptides from porcine teschovirus-1 (P2A: ATNFSLLKQAGDVEENPGP) and from Thoseaasigna virus (T2A: EGRGSLLTCGDVEENPGP) (**Figure 1B**).

### Proving sensor expression efficiency from CASPAM

Once CASPAM was generated, we set out to address the hydrolytic efficiency for the viral P2A and T2A sequences by examining the resulting cleavage products obtained upon transfection. For that purpose, HeLa cells were transiently transfected with CASPAM expression plasmid, harvested 24h later and subjected to western-blotting. As shown in **Figure 2A**, no faint bands corresponding to byproducts from an impaired or incomplete cleavage process were detected. Furthermore, fusion proteins consisting neither in the three biosensor moieties (*∼*170 kDa) nor two successive moieties (*∼*110 kDa for sCas3-GP-linker-P2A-sCas8-BP or sCas8-BP-linker-T2A-sCas9-RP) were observed. By contrast, a band with the expected molecular weight for a fluorescent protein dimer (*∼* 52 kDa) was the only identified product, additionally showing that no cleaved sensor is present in unperturbed conditions (**Figure 2A**). However, as anti-GFP antibody equally recognizes both Cerulean and mCitrine fluorescent proteins (but neither mCherry nor mKate), it was not possible by this means to fully determine successful biosensor co-expression. In this manner, although the existence of non-separated product due to failed 2A-linker cleavage was dismissed, it remained to be shown if the three biosensors were being simultaneously expressed in single cells. Thus, we set out to determine whether CASPAM enables co-imaging of the three biosensors.

**Figure 2:**
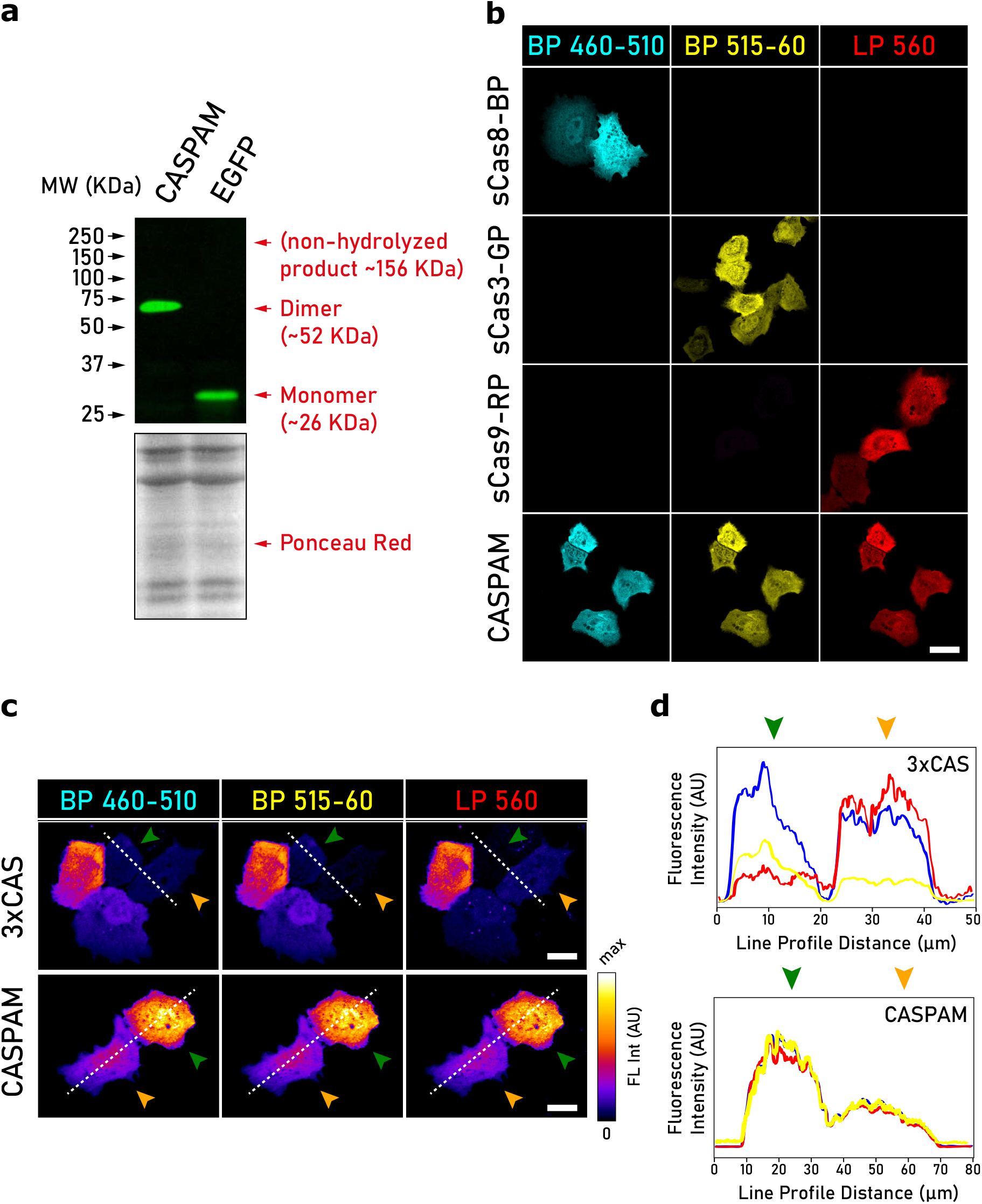
P2A and T2A sequences undergo successful hydrolysis, resulting in the effective simultaneous biosensor expression in cultured mammalian cells. **(a)** WB analysis of hydrolysis efficiency for the viral 2A sequences used in this study. Hela cells were transfected with the indicated plasmids and processed for WB 24h later. For the hydrolyzed product assessment, an anti-GFP antibody was used. A band matching with the expected molecular weight for a fluorescent protein dimer (*∼*52KDa) was the only product observed in CASPAM transfected cells. A lane corresponding to EGFP transfected cells is shown as a molecular weight reference control for a fluorescent protein monomer (*∼* 26KDa). Ponceau-red staining was used as a loading control. **(b)** Confocal microscopy imaging of the constructs used in this study. Hela cells were independently transfected with sCas3-GP, sCas9-BP, sCas8-RP or CASPAM expression plasmids. 24h after transfection, cells were fixed in 4% PFA (Materials & Methods) and subjected to laser scanning confocal microscopy imaging. Representative images of cells are shown. Scale bar: 10 *µ*m. **(c)** Confocal images of cells co-transfected with sCas3-GP, sCas9-BP and sCas8-RP expression plasmids (top) or CASPAM (bottom) as representative for the observed intercellular variability in fluorescence intensity. Colorbar indicates fluorescence intensity (FL Int) in arbitrary units (AU). Scale bar: 15 *µ*m. **(d)** Fluorescence intensity profiles of CASPAM (left) or sCas3-GP, sCas9-BP and sCas8-RP co-expressing cells (right). Fluorescence intensity plotted along the withe-dashed lines in panel **c** run through both the inside and the outside of the arrowed cells (blue: BP460-510; yellow: BP515-60; red: LP560).

Along with the aforementioned western-blot data, detection of fluorescence signal in the right emission channel would be indicative of proper expression and folding of each of the three genetically encoded biosensors. For this, HeLa cells were independently transfected with sCas3-GP, sCas9-BP, sCas8-RP or CASPAM plasmids. 24h after transfection, cells were PFA fixed and imaged with laser scanning confocal microscopy. Different detection channels were configured to image each individual biosensor. Due to the lack of significant spectral overlap between fluorophores [12] linear unmixing adjustments were dismissed. As shown in **Figure 2B**, none of the cells transfected with the plasmids encoding the individual biosensors showed fluorescence signal in a detection channel other than that specified for each fluorophore-pair. On the other hand, CASPAM transfected cells displayed simultaneous fluorescence signal in all three detection channels. These findings support both proper cleavage, expression and folding of the single biosensors from CASPAM. It is noteworthy that, although fluorescence often shows substantial intercellular variability, there is not such appreciable variability between detection channels for CASPAM transfected cells when compared with cells triple-transfected with sCas3-GP+sCas9-BP+sCas8-RP expression plasmids (3xCAS) (**Figure 2C-D**), suggesting for the formers a bona-fide correlation in fluorescence intensity.

### Fluorescence intensity ratio from any pair of biosensors remains constant in CASPAM

Regardless of the protein co-expression strategy choice, relative expression levels largely vary between individual cells [26]. This fact represents an obvious disadvantage, as hardly having any control over the expression ratio of key components of a given biological pathway tends to hamper quantitative studies. To address this issue, we set out to evaluate the performance of CASPAM to co-express proteins reliably at a defined ratio in single cells. For that purpose, HeLa cells were triple or single transfected with sCas3-GP+sCas8-RP+sCas9-BP (3xCAS) or CASPAM expression plasmids, respectively, and subjected to multicolor flow-cytometry analysis.

As evidenced by 2D histograms (**Figure 3A**), single cell biosensor co-expression from separate plasmids (by ordinary mixing equal amounts of plasmid in the transfection procedure, 3xCAS), resulted in widely varying cell-to-cell expression ratios. Quantification of the fluorescence emission intensity from single cells in the BP450/50 (Green), BP530/30 (Blue) and BP610/20 (Red) detection channels showed marked heterogeneity in the green to blue (GP:BP), green to red (GP:RP) and red to blue (RP:BP) expression ratios. Furthermore, sub-populations of cells expressing only one (or two) of the three biosensors were easily identifiable within 3xCAS transfected cells (**Figure 3A**). Conversely, in the cells transfected with CASPAM, co-expression takes place at a defined ratio, remaining virtually constant within the entire dynamic range (**Figure 3B**).

**Figure 3:**
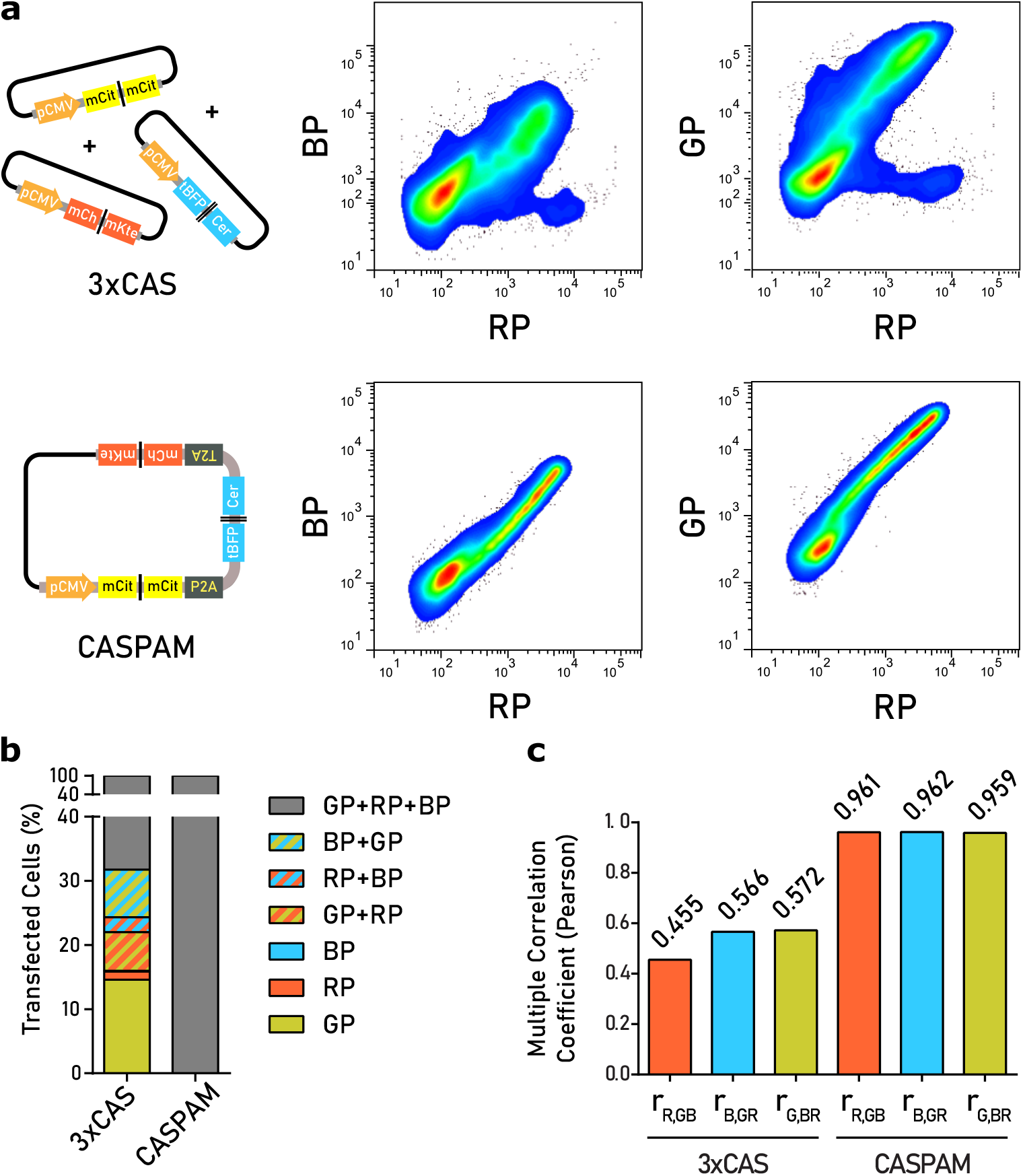
Comparison of co-expression strategies through the assessment of single cell sCasx-BP, sCasx-GP and sCasx-RP fluorescence intensity by multicolor flow-cytometry. Two different strategies for co-expression are analyzed: adding equal amounts of each biosensor-expression-plasmids in the transfection mix (3xCAS) versus CASPAM. **(a)** Bidimensional histograms obtained from flow-cytometry dot plots show fluorescence of mCitrine versus tagBFP/Cerulean, mCitrine versus mCherry/mKate and tagBFP/Cerulean versus mCherry/mKate, respectively (in arbitrary units). **(b)** Stacked-bar diagram showing the percentage of cells expressing one (plain colored bars), two (stripped bars) or three (dark grey bars) biosensors upon co-transfection with sCas3-GP, sCas9-BP and sCas8-RP (3xCAS) in comparison to CASPAM transfected cells. **(c)** For each co-expression approach, cell populations successfully expressing the three biosensors, were gate selected and subjected to Pearson multiple correlation analysis.

Although transfection efficiency can be experimentally tuned up, when cells were transfected with a mixture of different plasmids (3xCAS), at best 68,25% of transfected cells expressed all three biosensors, while 15,77% expressed two of them, and the remaining 15,97% a single-one. On the other hand, the entire pool of cells transfected with CASPAM expressed the three biosensors (**Figure 3C**). Moreover, the correlation of the fluorescence intensity ratios was significantly higher in CASPAM transfected cells, indicating that the variance in relative expression is smaller. Pearson multiple correlation analysis yielded coefficient values of r_*R,GB*_=0.455, r_*B,GR*_=0.566 and r_*G,BR*_=0.572 for the 3xCAS transfection, while for CASPAM, the values were r_*R,GB*_=0.962, r_*B,GR*_=0.962 and r_*G,BR*_=0.959, which showed a more linear-like correlation (**Figure 3D**). To sum up, CASPAM not only achieves successful co-expression of all biosensors, but also a better conserved co-expression ratio.

### CASPAM enables close to equimolar biosensor expression

A constant ratio in the fluorescence intensity values between a given pair of biosensors indicates a conserved ratio in the expression levels, but not an equal number of molecules, mainly due to differences in molecular brightness and acquisition parameters. Therefore we sought to determine the actual stoichiometry of the three biosensors in living cells using Fluorescence correlation spectroscopy (FCS). FCS is a confocal microscopy based, single molecule detection technique that measures and correlates fluctuations in intensity due to fluorescently labeled particles entering or leaving a femtoliter-scale observation volume [30, 31, 32].

HeLa cells were then transfected with either 3xCAS or CASPAM expression plasmids, and allowed to grow for 24h prior to FCS analysis, as detailed in Materials & Methods. In short, cells expressing all three biosensors were selected from each transfection condition, and fluorescence intensity fluctuations were measured in 10 repetitions of 10 second long, for each channel, in a single point per cell (**Figure 4A**).

**Figure 4:**
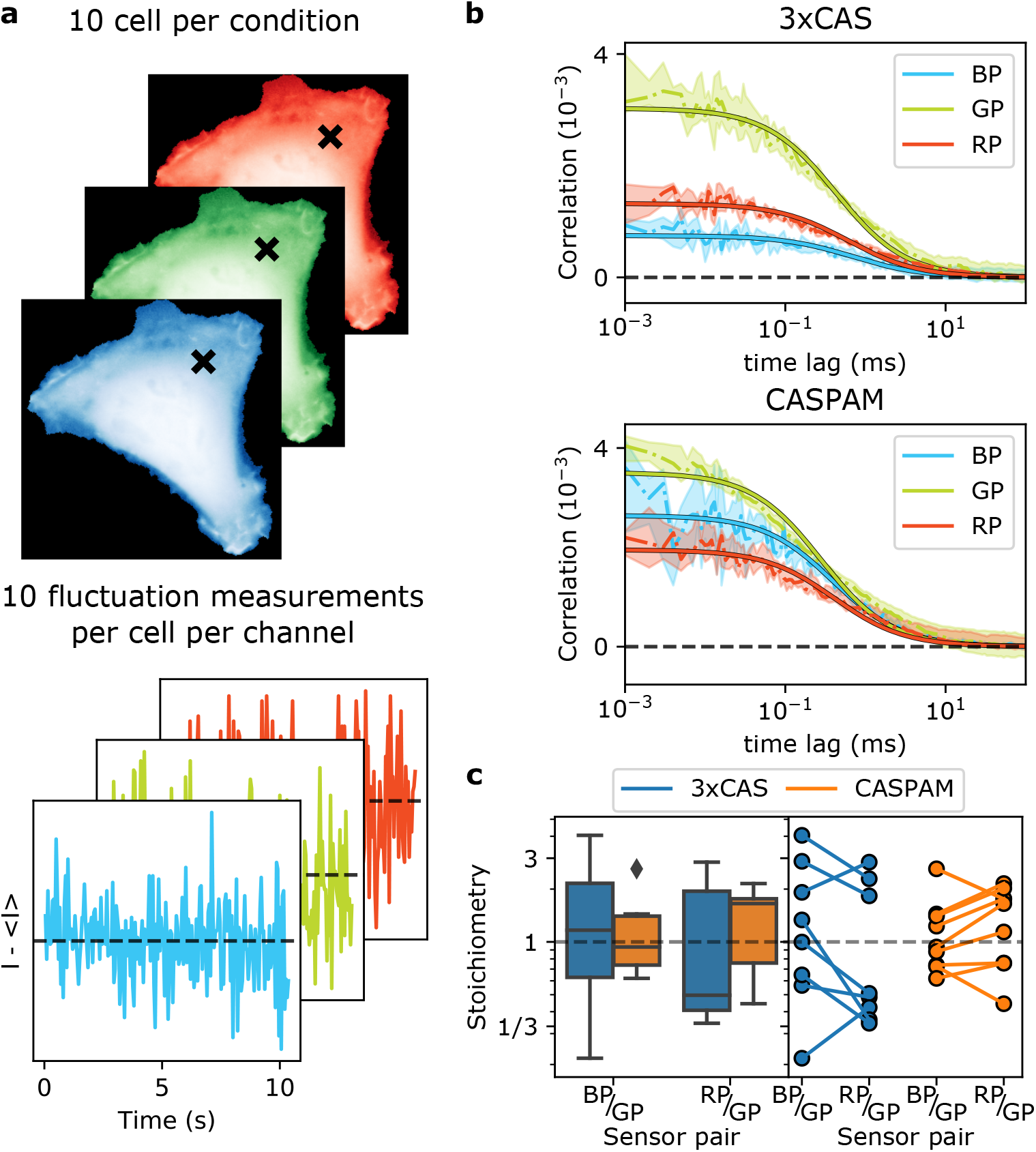
CASPAM enables near equimolar biosensor expression. **(a)** Schematic representation of a standard point scan fluorescence correlation spectroscopy (point FCS) pipeline. Fluorescence fluctuations of mobile fluorescently labeled proteins are acquired from a chosen location within an individual cell. 10 fluctuations were acquired per channel per cell, for 10 cells for either of the following conditions: three single plasmid and CASPAM transfection. **(b)** Operations on fluorescence signals, obtained by continuous scanning of a single pixel on the screen (the confocal volume), are carried out in point FCS by an autocorrelation function (ACF). Fitting the ACF to an appropriate model function results in obtaining concentration of several species and their diffusion coefficients. **(c)** The stoichiometry between biosensors was calculated obtaining for single plasmid per sensor a stoichiometry between BP and GP of 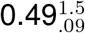 and for RP and 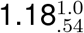 while for CASPAM the stoichiometry between BP and GP was 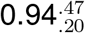 and between RP and 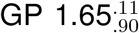 (where values represent the median and the interval between 25 and 75 percentiles). Triple-modality-reporter construct shows a relative expression between biosensors that is nearer to equimolarity (left). Paired plot of stoichiometry shows a reduced variability for CASPAM transfected cells (right).

As shown in **Figure 4B**, we calculated and fitted the autocorrelation function (ACF) for every cell and obtained both the concentration and the diffusion coefficient for each biosensor. As biosensors consist of dimers of similar size and shape, the diffusion coefficient was indistinguishable between cells, fluorescent protein pairs and transfection condition selected. Nonetheless, concentration of biosensors differed both between cells as well as between species inside each cell. In spite of the selection bias introduced by the fact that fluorescence microscopy and FCS have a narrow dynamic range in terms of fluorescence intensity, as compared to Flow Cytometry, the 3xCAS transfected cells showed a larger relative expression variance than CASPAM (**Figure 4C**). In particular, for 3xCAS, the stoichiometry between BP and GP remained within 0.41-1.95, and 0.63-2.16 for RP and GP, while for the CASPAM construct the stoichiometry between BP and GP was 0.74-1.40 and 0.76-1.76 between RP and GP (where values represent the median and the interval between 25 and 75 percentiles). Additionally, the relative expression of each biosensor pair for CASPAM proved to be near equimolar: 0.93 for BP and GP and 1.65 for RP and GP.

Taken together, these data support that CASPAM can successfully be used for tightly controlled, defined-ratiometric FPA caspase-activity biosensor co-expression in individual cells.

### Biological observables have less variability when perturbations are better controlled

Having successfully validated CASPAM, we set out to test it in a biologically relevant experimental setting: the induction of apoptosis by staurosporine (STS) [12, 33, 17, 34]. For this, HeLa cells, transfected either with the same amounts of plasmid for each biosensor (3xCAS) or with CASPAM were treated with STS and imaged for 15 hours. Acquired images were analyzed by means of a custom-made Python pipeline as described in Materials and Methods. As illustrated in Figure 5A (top row), no substantial variations in fluorescence intensity were observed. Consecutively acquired images for the fluorescence of the BFP channel, corresponding to a cell undergoing apoptosis, are shown as representative. However, changes in fluorescence polarization anisotropy were easily identifiable, and different for each of the three documented channels (**Figure 5A**-other rows). Anisotropy curves, as shown in **Figure 5B** (and **Supplementary Video V1**), were generated both for each channel and each segmented cell automatically, and these were further analyzed to obtain the time of maximum caspase activity (**Figure 5C**) as previously described [12].

**Figure 5:**
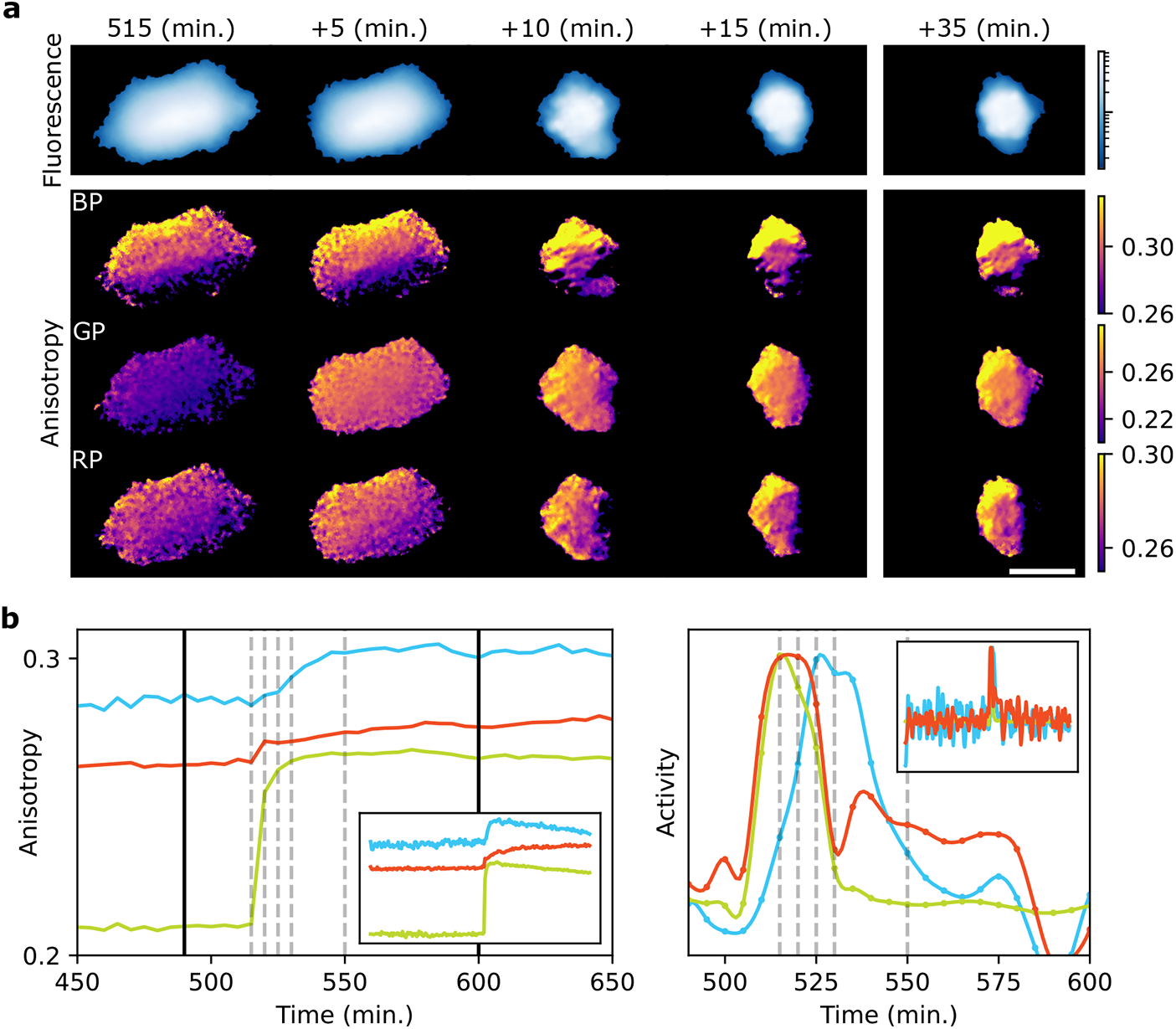
Characterization of fluorescence polarization anisotropy sensors for monitoring caspase activation. HeLa cells were stimulated with 1 *µ*M staurosporine. The caspase activity sensor cleavage was examined by quantification of the homo-FRET efficiency by fluorescence polarization anisotropy microscopy. **(a)** Consecutive images corresponding to blue channel fluorescence intensity and anisotropy images for each channel of a cell undergoing apoptosis. Differences in timing of cleavage of each sensor can be seen. Scale bar: 10 *µ*m. **(b)** The anisotropy curve of the corresponding cell is shown and dashed vertical grey lines correspond to each column of images (left). The instantaneous activity of each caspase is plotted (right). Estimated points are shown with dots and interpolated with splines. The difference in profile and time delay between caspase activities can be appreciated.

To address the impact of intercellular variability due to different concentrations between biosensors, we compared simulations with resampled experimental data. By means of an ordinary differential equation-based model previously introduced [12], we first simulated cells with varying concentrations of sCas3 and found that signal propagation is delayed with increasing concentrations (**Figure 6A**) as well as time differences of maximum activity between caspases (**Figure 6B**), as expected. For the experimental counterpart, we measured the impact of narrowing the concentration range on both the 3xCAS and CASPAM dataset. Cells whose intensity z-score was between -0.5 and 0.5 for each fluorescence channel separately (**Figure 6C**) were selected to emulate a narrower concentration range. After analyzing time differences of maximum activity between caspases, we found that standard deviation of time differences of each population was lower for cells transfected with CASPAM as expected (each row in Figure 6D). Indeed, as expression of different CASPAM FP are strongly correlated, computationally selecting an intensity range in one of them also selects an equivalent intensity range in the other two.

**Figure 6:**
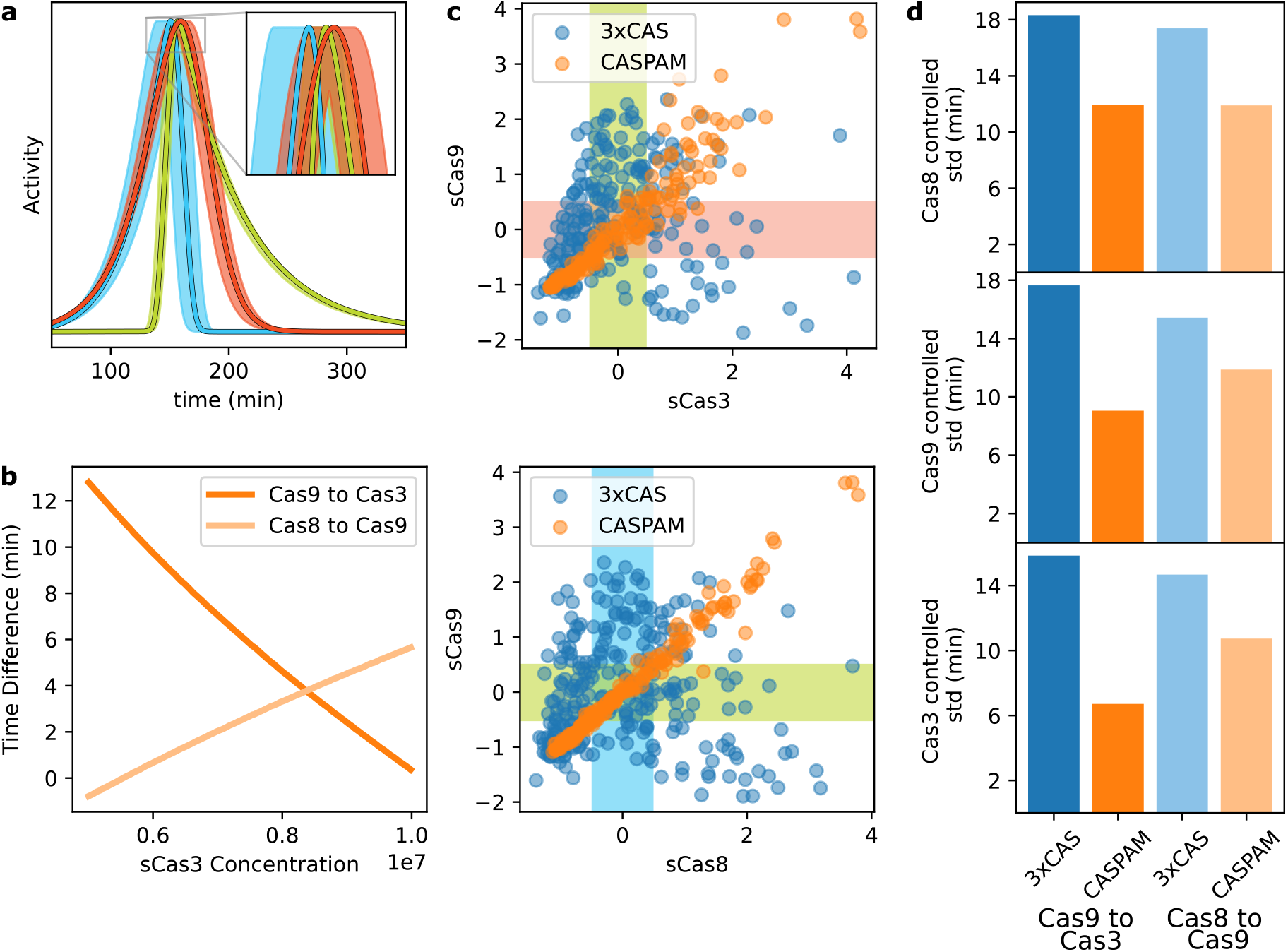
Uncorrelated expression leads to different delays in signal propagation throughout the network. **(a)** Simulated normalized activity profiles of a cell stimulated with an extrinsic ligand, using the model described elsewhere [12]. The filled region corresponds to simulated cells with different initial concentration of sCas3-FP. Plot with dark edges correspond to simulation with median concentration. **(b)** Time difference between maximum activity of caspase 9 and 3 and between caspase 8 and 9 was calculated and plotted against the initial concentration of sCas3-FP used in the simulation. **(c)** HeLa cells were triple transfected with a single plasmid per reporter (blue) and with CASPAM (orange). Scatter plot showing the dispersion of z-score of median fluorescence per cell of one biosensor against the other. A considerable improvement can be appreciated when CASPAM is used as values are much less dispersed. Boxes show cells selected whose total fluorescence z-score for a specific biosensor lies between -0.5 and 0.5. **(d)** Each row corresponds to selected cells according to intensity of a different channel. For each transfection condition, the standard deviation of time difference of maximum activity between caspases was calculated. It can be appreciated that CASPAM has less variability in time differences.

All in all, by using CASPAM and thus achieving near equimolar stoichiometry in expressed FPA caspase activity biosensors, perturbations to the system are much better controlled, leading to substantially less variation in biological observables. Furthermore, accurately knowing the relative concentration between biosensors allows for easier modelling as there is no need to estimate the range of concentrations of each introduced biosensor separately.

## Discussion

In this work we report the development of CASPAM, a single-plasmid-based triple-reporter-system that enables simultaneous imaging of the activity dynamics of three key nodes of the caspase network. These genetically-encoded biosensors can be monitored at single cell level for long extensions of time. Contrarily to more ubiquitous hetero-FRET biosensors, homo-FRET based biosensors take advantage of another light property, polarization as opposed to color, leading to a narrower spectral bandwidth per biosensor and the possibility of simultaneously imaging more biosensors in the visible spectra. Experimental multiplexing at the single cell level allows quantifying covariations in signals that could have been missed out by computational means and cope with intercellular variation in the process.

We characterized its functionality and successful co-expression capabilities by using both molecular biology and fluorescence-based techniques. We demonstrated that our 2A system provides reliable co-expression of all three functional reporters encoded within the single plasmid. This approach avoids the use of alternative methods that, although highly effective when transfecting several plasmids (electroporation or microinjection) tend to compromise cell integrity [35]. For infection purposes, 2A sequences (roughly 20aa long) hold great potential over IRES elements, whose relatively large size often collides with limited viral packaging capability [27]. In addition, 2A usage simplifies the generation of stably expressing cells by diminishing the number of independent transfection events required.

Using fluorescence correlation spectroscopy we also demonstrated that for the triple modality sensor the subunits are expressed in equimolar concentrations. Therefore, CASPAM is extremely convenient for high-throughput approaches. In particular, in microscopy experiments, cells or fields of view could be chosen by assessing a single channel; furthermore, imaging parameters could be selected just from a few sample cells avoiding, i.e., saturation due to other out-of-range cells.

The genetically encoded homo-FRET based biosensors used in this study rely on fluorescent proteins with orthogonal excitation-emission wavelengths, which enables monitoring at the same time the activity of each of the three main caspases from the apoptotic network. By contrast to the so called *computational multiplexing* [1, 6], experimentally multiplexed imaging emerges as a major importance procedure to overcome cell-to-cell and intracellular variability, with great potential for analyzing signaling hierarchies pathway kinetics in individual cells. Such capability is crucial in cases where a third node conditions the response of a pathway to some particular signal in a context dependent manner [33, 36].

Experimental restrictions for multiplexing approaches arise when attempting to increase the number of reporter molecules loaded into one cell. Apart from the limited tolerance to exogenously added factors, these latter represent a non-desirable perturbation of living cell homeostasis: titration and/or competition with endogenous components may perturb cellular response [12]. Thus, information regarding the perturbation introduced is valuable either for interpreting the data as for modeling purposes in reduction of parameters needed. Moreover, three is the minimum number of variables to do causal modelling [37], as we would like to detect (vanishing) conditional independencies, that is, explaining correlations between two variables as caused by a third. The strategy presented in this paper makes it possible to integrate into a pathway model, the heterogeneous and complex responses made visible by simultaneous imaging of different biosensors.

In summary, our molecular tool will provide a reliable methodology for accurately distinguishing between different caspase dependent biological landscapes: from signal propagation in the apoptotic network [12, 17], to non-apoptotic caspase activity, like dendrite pruning [38], and different types of apoptosis where not all caspases are active [33, 36], amongst others. Furthermore, the expressed biosensors allow refinement of existing models of the apoptotic signaling cascade. Applying different stimuli and perturbations to the network will provide new information of how signals are propagated throughout the network. What is more, perturbations introduced by CASPAM are better controlled for and thus, more precise modelling of the network is attainable. Concurrently, the implementation of this biosensor will make room for quantitatively studying the many biomedical processes where either apoptosis or non-apoptotic caspase activation occur in a spatially organized manner.

## Materials and Methods

### Cell culture & conditions

HeLa cells (ATCC, CCL-2) were purchased from American Type Culture Collection (ATCC, Manassas, VA, USA). Cells were cultured in high-glucose DMEM (Gibco) containing 10% FBS (Gibco) and supplemented with 10 mg/mL sodium pyruvate and 2 mmol/L L-glutamine plus antibiotics (100 U/mL penicillin, 100 *µ*g/mL streptomycin), and grown at 37 *°*C in a humidified incubator under a 5% CO_2_–95% air atmosphere. The day before transfection, cells were seeded in 8 well dishes (LabTekII, Nalgene) at a density of 3 *×* 10^4^ cells/well. Transient transfections were carried out using Lipofectamine 2000 (Invitrogen) according to the manufacturer’s instructions. 20-24h post transfection cells were imaged in DMEM without Phenol Red (PAN Biotech) and 0% FBS either in presence of 1 *µ*M Staurosporin (STS, Sigma Aldrich, Germany), to induce apoptosis, or DMSO, as a control, together with 10 *µ*g/ml Cycloheximide (Sigma Aldrich, Germany).

### Plasmid construction

Fluorescence polarization anisotropy caspase activity sensors were constructed as follows: a second fluorophore amplified by PCR and flanked by a SpeI and a SalI restriction site and containing a STOP codon was inserted into the SpeI/SalI restriction sites of a C1-vector backbone (Clontech). Additionally, the PCR product contained a short linker sequence at the 5’ end. These plasmids were used as non-cleavable controls. To construct the cleavable caspase sensors, the sequences encoding the cleavage sites of caspase 3, 8 and 9 (DEVD, IETD-IETD, LEHD, respectively) were amplified by PCR and the products, flanked by a BspI and a SpeI cleavage site, subcloned into the corresponding restriction sites of the digested non-cleavable control sensors. Primer sequences were previously described [12].

For the triple modality reporter construct, a single synthetic gene containing the aforementioned individual caspase activity sensors was designed (GenScript, Piscataway, USA). Briefly, the three sensors were arranged in tandem (see scheme in **Figure 1B**), with flexible Gly-Ser-Gly-linkers (GGGSGGG) [24] and viral 2A sequences (P2A: ATNFSLLKQAGDVEENPGP and T2A: EGRGSLLTCGDVEENPGP) between them. With exception of the fluorophore at the 3’ end of the construct, all native STOP codons were replaced by Gly. Finally, the synthetic gene was cloned within the KpnI/EcoRI restriction sites of the pcDNA3.1(+) vector (Life Technologies, Grand Island, NY). The integrity of each cassette in the mCitrine-DEVD-mCitrine-P2A-tagBFP-IETD-Cerulean-T2A-mCherry-LEHD-mKate triple modality reporter was confirmed by sequence analysis in the final construct.

### Western Blotting

Western blot was performed by standard procedures. Briefly, the day before transfection, Hela cells were seeded in 35mm dishes (2.5 *×*10^5^ cells/dish) and cultured in high-glucose DMEM in the conditions described above. Cells were either transfected with EGFP or with the triple-modality-reporter. 24h after transfection, samples were resuspended in Laemmli lysis buffer and boiled at 100 *°*C for 5 min. Protein samples were separated on a 12.5% SDS-PAGE gel and transferred to a polyvinylidenedifluoride (PVDF) membrane (Immobilon-P, Millipore). Blots were blocked with 3% non-fat milk-0.05% Tween-20 in Tris buffer saline (TTBS) and incubated with primary antibody anti-GFP (rabbit polyclonal, 1:1000 Santa Cruz Biotechnologies). After wash-out, incubation with secondary antibodies and Odyssey (LI-COR Biosciences, USA) mediated detection were performed according to manufacturer’s instructions.

### Confocal Microscopy of fixed cells

Hela cells grown as stated in Cell culture & conditions were fixed in 4% paraformaldehyde, 4% sucrose, and 2 mM MgCl_2_ in PBS for 20 min at room temperature. Samples were mounted in MOWIOL 4-88 (Sigma Aldrich) and images were acquired by a confocal scanning microscope (Spectral FV1000 Olympus) using a 60X UPLSAPO oil immersion objective lens with a numerical aperture (NA) of 1.35.

### FCS measurements

Hela cells were seeded in 8 well dishes (LabTekII, Nalgene) in the conditions previously described (Cell culture & conditions) and transfected with i) a DNA mix containing equal amounts of sCas3-GP (mCitrine-DEVD-mCitrine), sCas9-BP (tBFP-IETD-Cerulean) and sCas8-RP (mCherry-LEHD-mKate) plasmids or ii) with the mCitrine-DEVD-mCitrine-P2A-tagBFP-IETD-Cerulean-T2A-mCherry-LEHD-mKate triple modality reporter construct. 20-24h after transfection, LabTeks were transferred to a Leica TCS SP8 inverted confocal scanning microscope (Leica Microsystems) equipped with a pulsed 470-670 nm white light laser (WLL2 Kit, NKT Photonics), an environment-controlled chamber (Life Imaging Services) and maintained at 37 *°*C in DMEM (with 25mM HEPES, without Phenol Red). A HC PL APO 63x/1.4NA CS2 oil objective (Leica Microsystems) was used. Fluorescence intensity fluctuations were collected by point-scanning in 10 repetitions of 10 seconds and corresponding ACF (auto-correlation function) curves were fitted to this data.

### Multicolor flow-cytometry measurements

Hela cells were seeded in 100-mm Petri dishes (1 *×* 10^6^ cells/dish) and cultured for 24h in the conditions previously described (Cell culture & conditions). For the proper flow cytometry settings adjustment, cells were transfected as follows: i) sCas3-GP, ii) sCas8-RP, iii) sCas9-BP, iv) sCas3-GP + sCas8-RP, v) sCas3-GP + sCas9-BP, vi) sCas8-RP + sCas9-BP, vii) sCas3-GP + sCas8-RP + sCas9-BP and viii) mCit-DEVD-mCit-P2A-tBFP-IETD-Cer-T2A-mCh-LEHD-mKte (CASPAM). 24h after transfection, the culture medium from each dish was collected and cell monolayers were then washed two times in 5 ml PBS, trypsinized, gently resuspended in 3 ml of complete medium and immediately mixed with the previously collected culture medium. Cells were then centrifuged (200-400 g, 10 min) and fixed in 4% paraformaldehyde, 4% sucrose, and 2 mM MgCl_2_ in PBS for 20 min at room temperature. After fixation, cells were centrifuged (200-400 g, 10 min), resuspended in 0.5 ml PBS and stored at 4 *°*C. Flow cytometry measurements were performed on a FACS-Aria Fusion cell cytometer (BD Biosciences) equipped with 405nm, 488nm and 561nm lasers. For choosing the appropriate laser and filter set configurations for each biosensor and achieve a clear discrimination of cells expressing one, two, three biosensors, or neither, experiments involving different combinations of them were firstly performed (**Supplementary Figures S2-S3**). Fluorescence emissions were detected for mCitrine, tag-BFP/Cerulean and mCherry/mKate using band-pass filters 450/50, 530/30 and 610/20 respectively. For each sample, a total of 2 *×*10^4^ counts gated on a forward scatter (FSC) versus side scatter (SSC) dot plot -excluding doublets-were analyzed. Data was acquired using CellQuest software (BD Biosciences) and processed with FlowJoTM v7.6.5 software. Pearson multiple correlation analysis was calculated.

### Fluorescence polarization anisotropy (FPA) microscopy

Data for anisotropy imaging was acquired with a custom-built setup, as previously described [12]. Briefly, images were acquired using an Olympus IX81 inverted microscope (Olympus, Germany) equipped with a MT20 illumination system. A linear dichroic polarizer (Meadowlark Optics, Frederick, Colorado, US) was placed in the illumination path of the microscope, and two identical polarizers were placed in an external filter wheel at orientations parallel and perpendicular to the polarization of the excitation light. Fluorescence was collected via a 20X NA air objective, and parallel and perpendicular polarized emission images were acquired sequentially on an Orca CCD camera (Hamamatsu Photonics, Japan). The CellR software (Olympus, Germany) or an in-house developed LabVIEW (National Instruments) program was used to control the data acquisition. Before each experiment, a reference sample consisting of a dilute fluorescein solution was measured. The samples were imaged at 37 *°*C using a temperature control system consisting of an objective heater and the Stable Z specimen warmer (Bioptechs Inc., Butler, PA, USA). Images were acquired every 10 min for a period of 15 h in the experiment for transfection with three different plasmids or every 5 min for 12 h in the experiment where all three sensors were transfected in a a single construct. Staurosporine (STS) was added to the wells in order to achieve 1 *µ*M concentration immediately before imaging.

### Image processing and analysis

mCitrine channel was used to generate segmentation masks of each cell by applying methods from the Cellment package (https://github.com/maurosilber/cellment). First, background pixels were identified by means of the SMO operator and after determining background pixels, a plane was fitted and subtracted from the image in order to subtract the background from every image. Remaining pixels were thresholded to be above the 70 percentile and morphological operations were applied to smooth borders and remove salt and pepper effect. Taking advantage of the fact that apoptotic cells usually detach from the slide and surrounding cells, a tracking method from the Cellment package was used to separate cells that are touching each other. After segmentation, total intensity for each channel at each timepoint was estimated for each cell, from which total fluorescence and anisotropy was subsequently calculated. Masked objects that do not correspond to apoptotic cells were filtered out, caspase activity curves were calculated for each apoptotic cell time lapse and time of maximum activity was obtained, as previously described [12].

### Modelling

We implemented the model introduced in Corbat et al., 2018 [12] and ran several simulations while sweeping concentration of sCas3 through expected values (0.5-1*×* 10^7^). Simulated results were analyzed in the same way as experimental ones.

## Supplementary

**Supplementary Figure 1:**
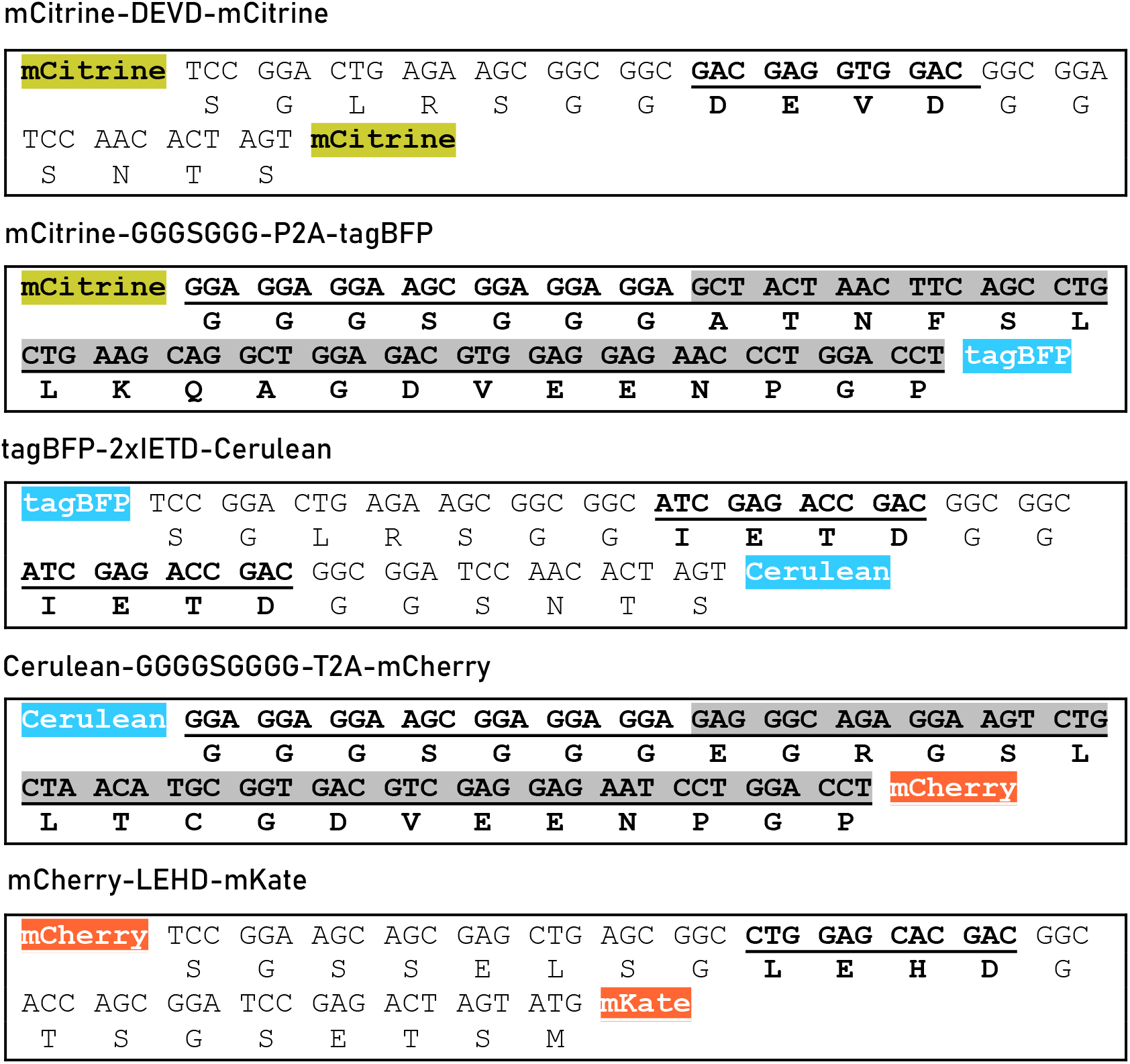
DNA and corresponding amino acid sequences of caspase cleavage sites and 2A peptides used in this study for mCitrine-DEVD-mCitrine, P2A, tagBFP-2xIETD-Cerulean, T2A and mCherry-LEHD-mKate respectively. Underlined sequences encode functional amino acids. For clarifying purposes, the four N-or C-terminal aminoacids of each fluorescent protein were included in each sequence.

**Supplementary Figure 2:**
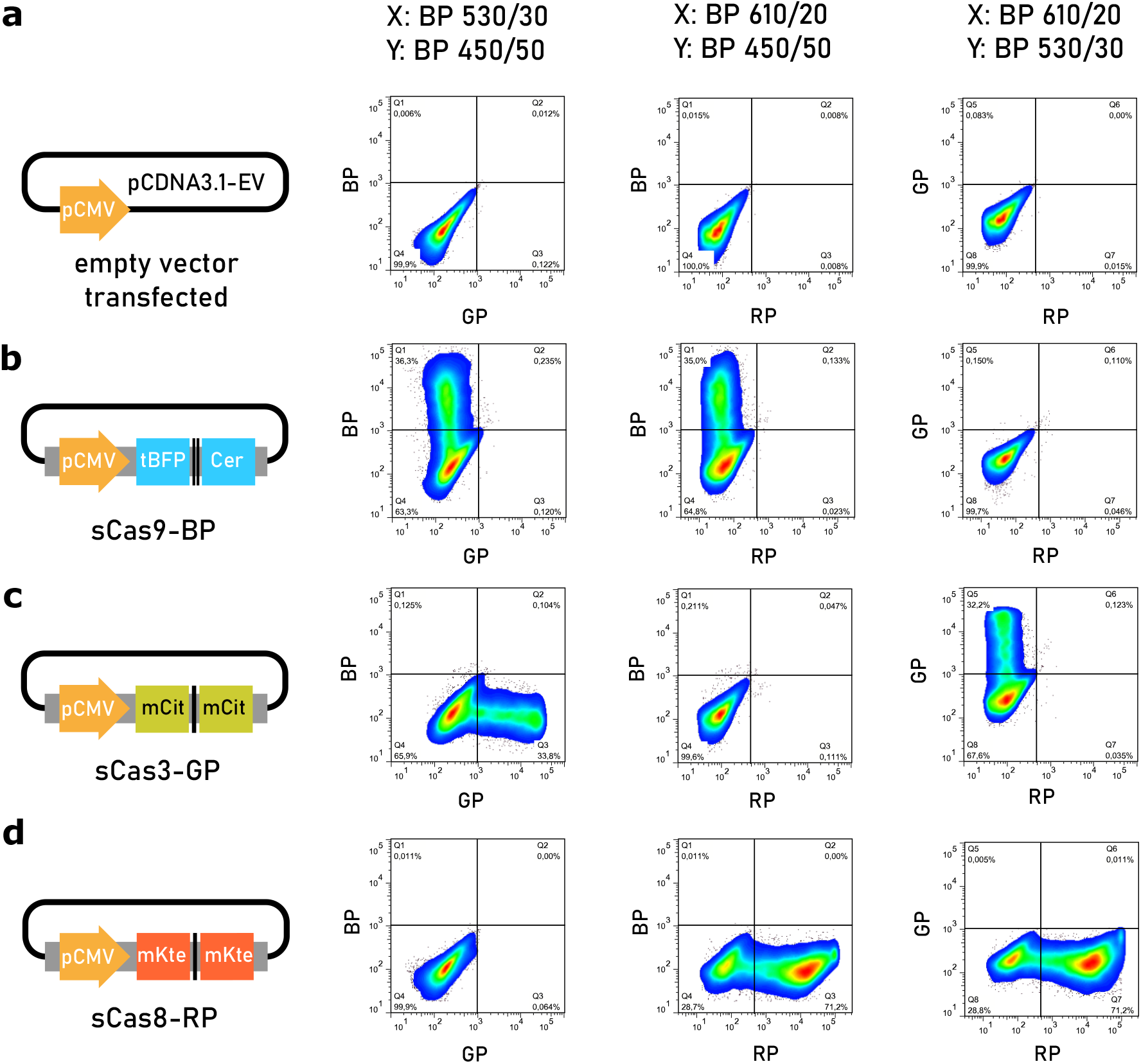
Flow cytometric detection of single biosensor expression. Hela cells were transfected with plasmids encoding a single FPA caspase activity biosensor, as detailed in Materials & Methods. Briefly, transfected cells were grown for 24h prior to harvest and analysis by flow cytometry. Samples were acquired using FACS-Aria Fusion cell cytometer, and a minimum of 20,000 events were collected for each sample. As seen from the representative histograms only two cell populations were detected: cells expressing one biosensor and untransfected/non biosensor-expressing cells (percentages are indicated on the plots). **(a)** pcDNA3.1 empty vector, **(b)** sCas9-BP, **(c)** sCas3-GP, **(d)** sCas8-RP. Single biosensor expression assessments served as single-color compensation controls to calculate the amount of compensation required for the co-expression determinations.

**Supplementary Figure 3:**
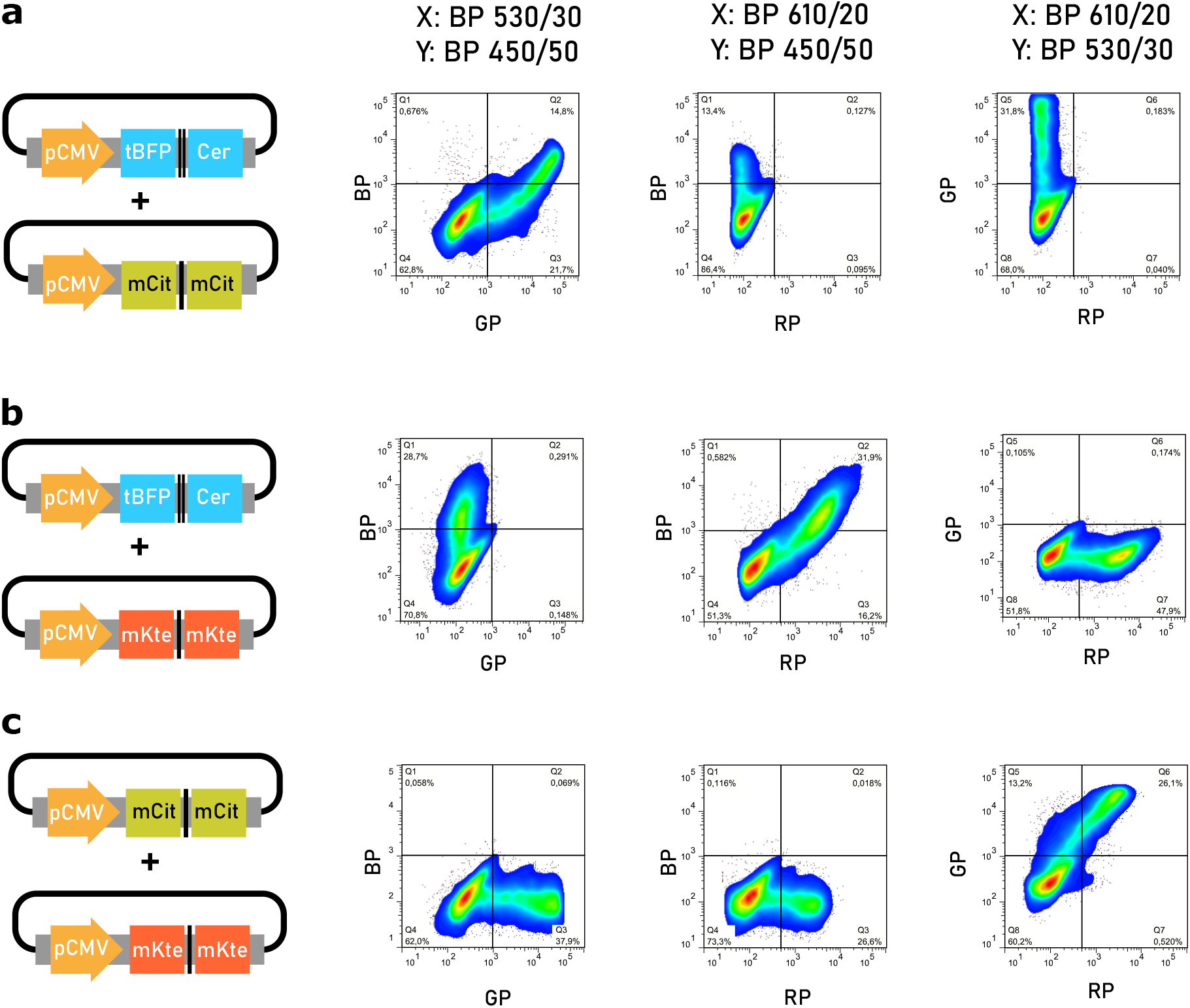
Flow cytometric detection of single cell dual biosensor co-expression. Hela cells were transfected with different combinations of two plasmids, each one encoding a single FPA caspase activity biosensor in 1:1 (w/w) mixes, as detailed in Materials & Methods. As seen from the representative histograms, all major cell populations were detected: cells expressing one of the biosensors, both biosensors, and neither biosensor (percentages are indicated on the plots). Cells expressing the biosensors are easily distinguished from those untransfected/non–biosensor-expressing cells. **(a)** sCas9-BP+sCas3-GP, **(b)** sCas9-BP+sCas8-RP, **(c)** sCas3-GP+sCas8-RP.

**Supplementary video V1.** Fluorescence intensity and anisotropy video corresponding to each channel of the cell presented in **Figure 5**. Anisotropy curve is also plotted below with a dashed line showing the actual frame.

## Acknowledgements

We thank Sabina Victoria Montero and Mariela Bollati-Fogolín from the Cell Biology Unit (UBC), Institut Pasteur de Montevideo, Uruguay, for their professional advice and technical assistance when performing the multicolor-flow cytometry determinations. We thank Dr. Sven Mü ller for technical assistance and Dr. Klaus Schuermann for useful discussions. We also thank Consejo Nacional de Investigaciones Científicas y Técnicas (CONICET) and University of Buenos Aires (UBA) for financial support to MH, AAC, MS and HEG. This work was supported by the following grants: PICT 2014-3658; PICT 2013-1301; MPG Partner Group.

## Author Contributions

MH, AAC, and HEG conceived the study and designed the experiments. MH designed the CASPAM construct and performed the biochemical, confocal microscopy, flow-cytometry and FCS-based characterization. HEG and AAC analyzed the FCS data, built the models and simulations and network analysis, and performed and analyzed the FPA microscopy experiments. MS designed the Cellment package for the segmentation of FPA images and analyzed the FPA data. MH, AAC, MS and HEG designed the figures and wrote the article. HEG proposed the idea of CASPAM and coordinated the whole project. All authors revised the manuscript and approved the submitted version

## Competing Interests statement

The authors declare that the research was conducted in the absence of any commercial or financial relationships that could be construed as a potential conflict of interest.

## Notes

### Competing Interest Statement

The authors have declared no competing interest.

